# Apparent Anatomical Variability Through Rigid Augmentation Enables Reliable Corpus Callosum Segmentation

**DOI:** 10.64898/2026.06.26.734817

**Authors:** Daniel M. Guimarães, Diego Szczupak, Vinícius P. Campos, Ivanei E. Bramati, Afonso C. Silva, Fernanda Tovar-Moll

**Affiliations:** D’Or Institute for Research and Education, Rio de Janeiro, Brazil; University of Pittsburgh, Pittsburgh, United States of America

## Abstract

The corpus callosum is a major white matter bundle responsible for connecting both hemispheres. In mammals, due to a variety of causes, the development of the corpus callosum can be impaired - this brain malformation is known as corpus callosum dysgenesis (CCD). The clinical presentation of CCD varies, with patients exhibiting three morphological phenotypes: agenesis, partial dysgenesis, and hypoplasia. Although the first two presentations are easily detectable on MRI scans, the latter is more challenging, as the structure is fully formed but has a reduced area. In this study, we develop (1) a pipeline to generate synthetic MRI scans with apparent anatomical variation and (2) train a U-Net–based tool to automatically segment the corpus callosum of marmosets in both healthy and disease contexts. Methodologically, a custom script was devised to apply rotation and translation to T1-weighted MRI scans at the volume level. Because the slicing grid remains unchanged, these rigid transformations translate into apparent anatomical variations at the slice level. We compared corpus callosum measurements obtained from automatically segmented masks with those from manually delineated masks. The average Dice score was above 0.90, and the Hausdorff distance was below 0.4 mm. We also stratified our cohort according to phenotype (healthy controls and hypoplastic animals). The magnitude of the effect and the significance level observed between the voxel counts of healthy and hypoplastic animals using manually delineated masks were comparable to those obtained via automatic segmentations. These results show that our pipeline can generate a sufficiently varied training pool to build an accurate U-Net segmentation model with high diagnostic capability.

## Introduction

Congenital brain malformations are heterogeneous, complex, and challenging to characterize (Brock et al., 2021; Severino et al., 2020; Thalhammer et al., 2025). These conditions may arise from a variety of causes, including environmental factors such as radiation exposure (Jayan et al., 2022) and gestational alcohol consumption (Jarmasz et al., 2017), as well as disruptions in fundamental neurodevelopmental processes such as axonal guidance and neuronal migration (Qian et al., 2022; Tsai et al., 2020), and glial development (Wang et al., 2022). They can manifest as diverse anatomical abnormalities, including aberrant cortical layer organization, e.g., subcortical band heterotopia, also known as double cortex syndrome (Chiba et al., 2022), complete absence of brain structures, such as in corpus callosum agenesis (Paul et al., 2007; Szczupak et al., 2021; Tovar-Moll et al., 2007), and incomplete encephalon growth due to cellular loss, e.g., microcephaly (Alcantara & O’Driscoll, 2014).

The detection of these anatomical abnormalities via imaging methods is routinely used in clinical practice to support both prenatal and postnatal diagnoses. Structural magnetic resonance imaging (MRI, a non-invasive technique with high spatial resolution, enables the identification of abnormal developmental landmarks from fetal stages throughout development such as altered head circumference (Hazlett et al., 2005; Rau et al., 2021; Selvanathan et al., 2022), as well as structural defects like schizencephaly (George et al., 2025). These imaging findings often prompt closer monitoring and referral to specialized care.

One of the most emblematic and prevalent developmental brain malformations is the corpus callosum dysgenesis (CCD) (Paul et al., 2007; Szczupak et al., 2021; Tovar-Moll et al., 2007, 2014). CCD encompasses a spectrum of abnormalities, including agenesis (aCCD), characterized by the complete absence of the corpus callosum; partial dysgenesis (pCCD), in which only a small portion - typically the genu - is formed; and hypoplasia (hCCD), in which the corpus callosum is fully formed but reduced in size. While aCCD and pCCD are often readily identified on neuroimaging, hCCD is characterized by a fully formed but reduced corpus callosum and may be overlooked or mistaken for a normal variant on qualitative inspection alone, particularly in borderline cases, emphasizing the need for quantitative morphometric assessment (Bodensteiner et al., 1994).

Quantification of parameters such as the area and volume of regions of interest (ROIs) often relies on manual delineation, making it susceptible to significant inter- and intra-observer variability (Kocak et al., 2019). Even when performed by expert anatomists, this process can introduce biases that affect the reliability of downstream analyses. Furthermore, purely qualitative assessments, in the absence of robust quantitative measures, may lead to false-positive or false-negative findings, ultimately compromising diagnostic accuracy (Dixon et al., 2022; Liu et al., 2024; Saltarelli et al., 2025). These limitations highlight the need for automated, objective, efficient, and reproducible tools for anatomical quantification.

Numerous deep learning-based algorithms have been developed to address brain segmentation tasks (Andrews & Ieva, 2025; Chau et al., 2025), both in healthy populations and in the presence of pathology (Koc & Akgun, 2026; Thabet et al., 2025). However, these models typically require large and diverse training datasets to learn robust and generalizable feature representations. Their performance is therefore strongly dependent on the quality and variability of the training data. In the context of rare conditions, where access to extensive imaging datasets is limited, this requirement becomes a major constraint. To mitigate this issue, image augmentation techniques have been proposed to artificially increase the variability of training sets (Chlap et al., 2021; Pani & Chawla, 2024).

Animal models, particularly rodents and non-human primates, play a central role in investigating neurobiological mechanisms underlying brain development and malformations (Edwards et al., 2020; Szczupak et al., 2023; Szczupak, et al., 2020a). However, differences in brain size, morphology, and image acquisition protocols pose additional challenges for automated segmentation methods (Despotović et al., 2015; Puzio et al., 2025; Rao et al., 2022). In this context, establishing reliable automated segmentation strategies in the marmoset brain represents an important step toward improving quantitative neuroanatomical analyses in translational research.

In this study, we propose a pipeline that leverages augmented MRI data to train U-Net-based models for the automatic segmentation of the corpus callosum in T1-weighted anatomical scans. We evaluate this approach using the marmoset as model and assess whether rigid data augmentation alone is sufficient to achieve accurate segmentation performance of the corpus callosum.

## Materials and Methods

### Sample characteristics

Our sample consisted of 45 adult marmosets (Callithrix jacchus): 38 with a normotypical corpus callosum (hereafter referred to as controls) and seven diagnosed with corpus callosum hypoplasia by an expert. Of the 38 control animals, eight were used to generate augmented scans, whereas the remaining 30 were used for model validation. The hypoplastic animals were not included in the training set and were used exclusively for model validation.

### MRI acquisition

MRI scans were acquired at the University of Pittsburgh using a 9.4T, 30 cm bore magnet (Bruker, Billerica, USA) equipped with a 20 cm gradient set capable of 300 mT/m (Resonance Research Inc., Billerica, USA). Radiofrequency transmission was performed using a 16-rung high-pass birdcage coil, and signal reception was achieved with a custom-built 16-channel phased-array coil.

High-resolution whole-brain 3D T1-weighted images were acquired using a Magnetization Prepared–Rapid Gradient Echo (MP-RAGE) sequence with the following parameters: inversion time (TI) = 1400 ms, echo time (TE) = 3.3 ms, repetition time (TR) = 6000 ms, flip angle = 12°, field of view (FOV) = 42 × 35 × 25 mm, and matrix size = 168 × 140 × 100. This configuration yielded an isotropic resolution of 250 µm with an acquisition time of approximately 20 minutes. A saturation band was positioned at the base of the marmoset’s head to suppress signal aliasing from regions outside the FOV.

### Imaging processing

Images were denoised using a variance stabilization transformation (VST) followed by the block-matching and 3D filtering (BM3D) algorithm. Noise in magnitude MR images is known to follow a Rician distribution (Foi 2011). To address this, we first applied the forward VST to stabilize the noise variance, converting signal-dependent noise into signal-independent noise with a Gaussian-like distribution. Subsequently, BM3D - a non-local transform-domain filter designed for volumetric data corrupted with Gaussian noise - was applied to denoise the images in the VST domain (Maggioni et al. 2012). Finally, the inverse VST was used to restore the data to its original magnitude space. No spatial resampling was performed, and all images were maintained at their native isotropic resolution (250 µm) throughout the processing pipeline.

### Setting up the environment

All scripts were implemented in Python and executed using Jupyter Notebooks within the Google Colaboratory environment. The models were run locally, with notebooks deployed using Docker (version 4.33.1). The hardware specifications of the local machine included 32 GB of RAM, an NVIDIA RTX 4060 GPU with 8 GB of dedicated memory, and an Intel Core i7-14700 CPU. Versions of all Python libraries used are documented within each script.

### Synthetic scan generation

Eight original T1-weighted scans underwent an image augmentation protocol based on two rigid motion transformations: rotation and translation. Rotation was performed using the sitk.Euler3DTransform function, while translation was applied using sitk.TranslationTransform. The image volumes were uniformly rotated by a randomly selected angle between –10° and +10°. Translation was fixed at 2 units. Both rotation and translation were applied along all three spatial dimensions (x, y, and z), and each transformation was followed by a resampling step.

For each original scan, five augmented images were generated. Although only three-dimensional rigid transformations were applied, apparent slice-level anatomical variability emerges in the augmented scans because the fixed sagittal slicing plane intersects the rotated volumes at slightly different anatomical angles after resampling and interpolation to the original voxel grid (Fig. 1).

**Figure 1.**
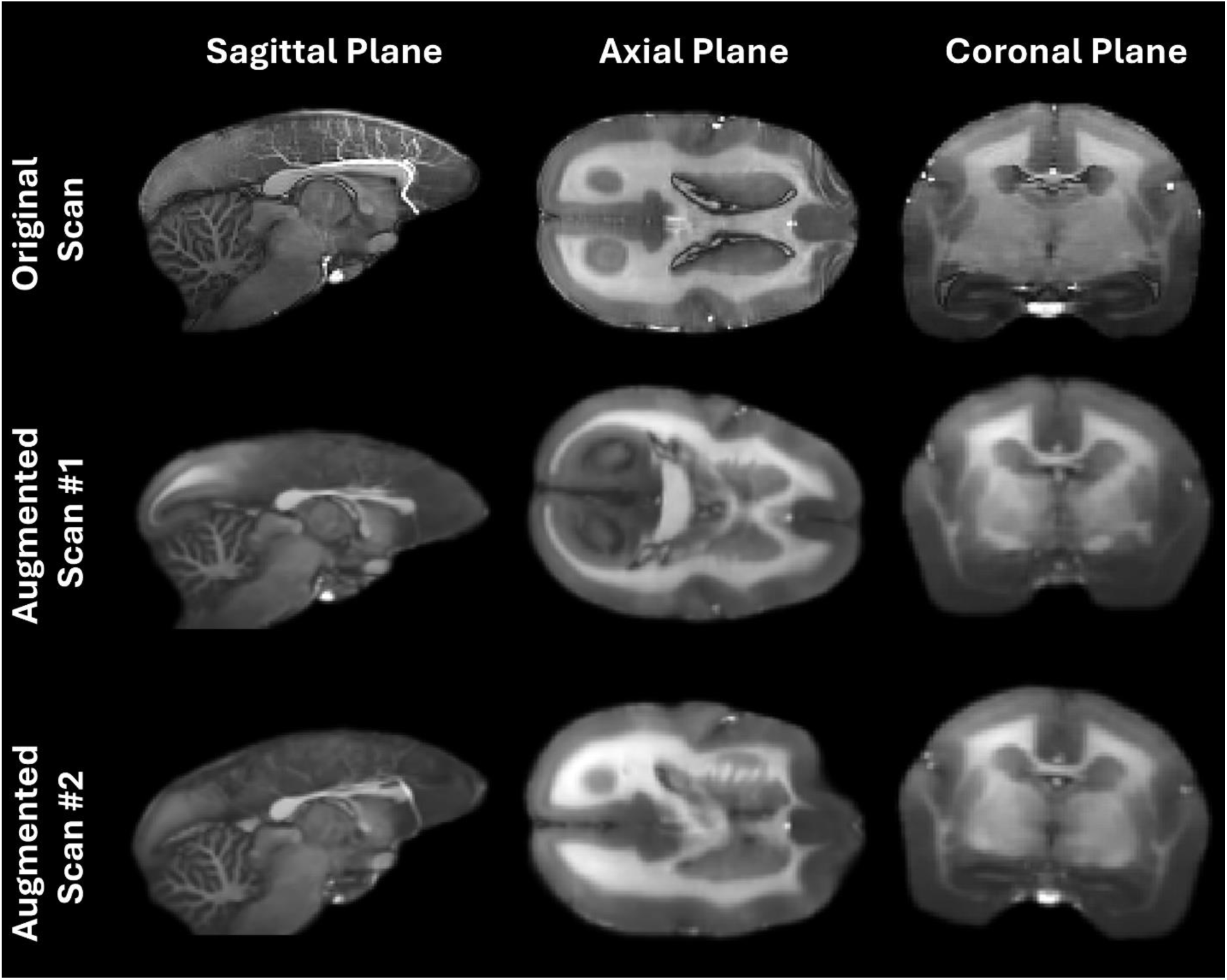
Examples of anatomical variations introduced in three augmented images derived from a single original scan. The images correspond to the same anatomical landmark (in the midline, where the fornix meets the corpus callosum) displayed in the three anatomical planes.

To quantify the differences, the mean absolute error (MAE) was calculated between the augmented images and their corresponding originals. Parameters for rotation (angle range) and translation (magnitude) were selected to ensure that augmented images were sufficiently distinct from both the original scans and each other, while maintaining anatomical plausibility. The Python script used for image augmentation will be made available upon request.

### U-Net segmentation training

Since the goal of the proposed pipeline is to train segmentation tools for diseases with low clinical representation, our deep learning model was trained exclusively with augmented images.

Forty augmented scans, along with their corresponding hand-annotated masks of the corpus callosum, were used as input to the model. The masks were manually drawn using the FSLeyes software (Jenkinson et al., 2012). For mask delineation, we considered the corpus callosum in the midline sagittal slice and parasagittal slices (one slice on each side or more depending on the morphology of the structure). To avoid measurement contamination, corpus callosum volume estimations were performed solely on the sagittal slices. In addition, both input images and their masks had their intensities normalized, and the slices dimensions were adjusted prior to the training.

The dataset was split into training and test sets using an 80/20 ratio, respectively. In addition to the volumetric rigid augmentations applied prior to slice extraction, further in-plane data augmentation was performed during training using Keras’ ImageDataGenerator. Whereas the initial transformations operate in the three-dimensional domain and induce apparent anatomical variability at the slice level due to fixed-plane slicing after rotation and resampling, the in-plane augmentations act on the two-dimensional slice representation, promoting robustness to spatial variations such as rotation (up to 40°), translation (width and height shifts up to 20%), scaling (zoom range of 20%), and shear (20%), with horizontal flipping enabled and nearest-neighbor filling. These two augmentation stages therefore operate in complementary geometric domains and are not redundant. The same transformations were applied simultaneously to MRI scans and their corresponding masks using a fixed random seed to ensure spatial correspondence. Further implementation details can be found in the scripts.

The network architecture was consistent across all three models: the encoder-decoder U-Net featured layers with filter sizes of 64, 128, 256, 512, 1024, followed by the decoder layers with 512, 256, 128, and 64 filters (Fig. 2). All convolutional layers used kernels of size 3, except for the 2D transposed convolutional layer which used a kernel of size 2. Rectified Linear Unit (ReLU) activations were applied at every layer except the final one, which used a sigmoid activation function. Batch normalization and dropout regularization were incorporated at every layer (except the last decoder layer) to reduce overfitting and prevent gradient explosion.

**Figure 2.**
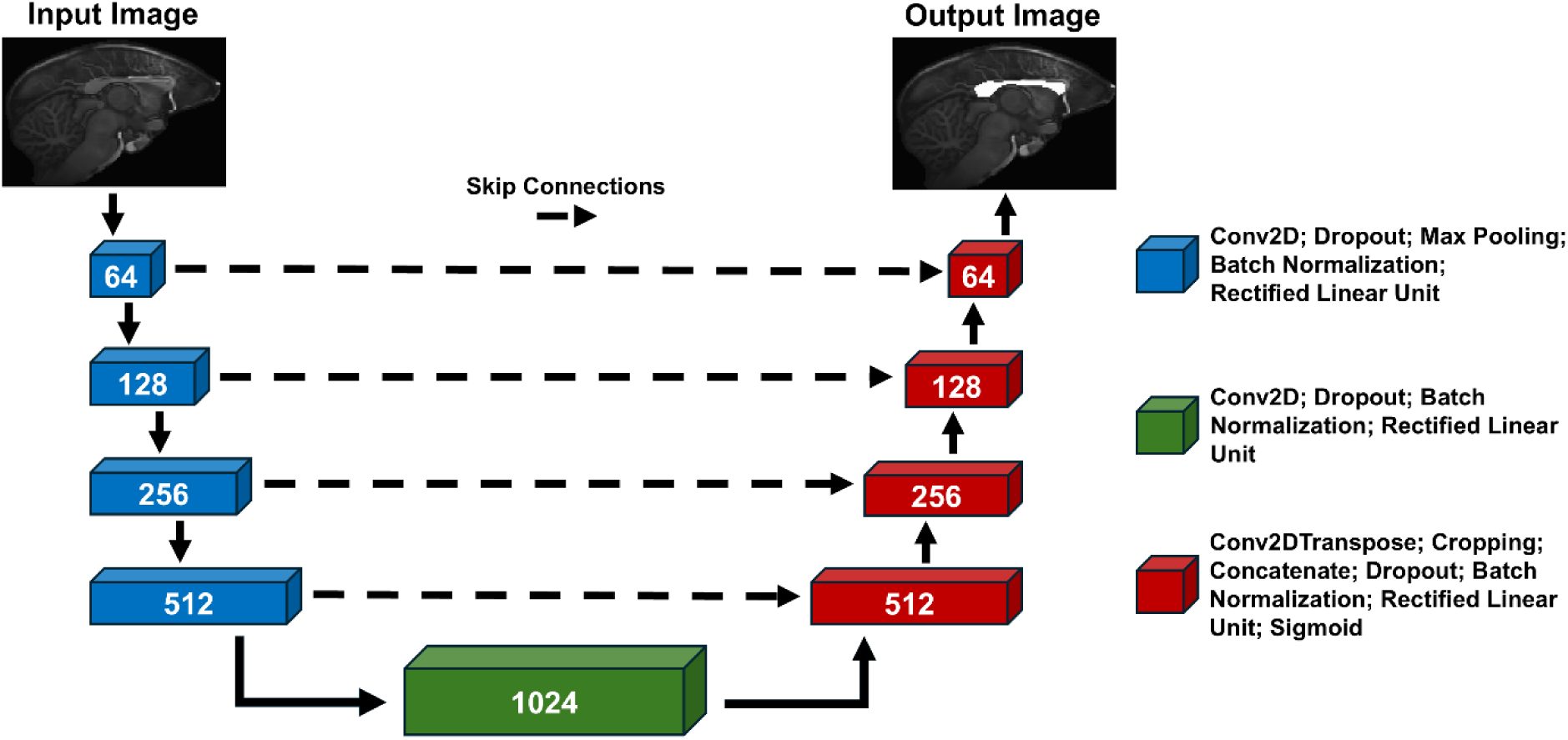
Schematic representation of the U-Net architecture used for corpus callosum segmentation. The encoder (left) extracts hierarchical features through successive convolutional blocks with downsampling, reaching a 1024-filter bottleneck. The decoder (right) reconstructs the segmentation map via transposed convolutions and upsampling. Skip connections transfer high-resolution features from encoder to decoder, preserving spatial detail for accurate mask prediction.

Training employed the Adam optimizer, with binary cross-entropy as the loss function. Two callback functions were used during training: a custom checkpoint to save the model state every *n* epochs, and ReduceLROnPlateau to decrease the learning rate if the loss did not improve after *n* epochs. Model selection was based on accuracy and loss metrics, and the best-performing model was then used to segment the corpus callosum in new scans. The best-performing model was trained for 3,200 epochs, achieving a final loss of 2.18 × 10⁻⁴ and an accuracy of 0.9998.

### Statistical analysis

Statistical analyses were designed to evaluate three complementary aspects of the proposed segmentation pipeline: (i) volumetric agreement between manual and automatic segmentations, (ii) spatial agreement between the resulting masks, and (iii) preservation of biologically meaningful volumetric differences between phenotypes. Importantly, all validation analyses were conducted exclusively on original scans that were not used during model training and were therefore unseen by the model.

To assess volumetric agreement between manual annotations and automatic segmentations of the same scans, two-tailed paired t-tests were performed. To evaluate volumetric differences between phenotypic groups (control vs. hypoplastic), two-tailed independent t-tests were used. Data are presented as scatterplots with paired observations connected by lines and group means indicated by red dots. Statistical significance was defined as p < 0.05.

Potential systematic biases between methods were further investigated using Bland-Altman plots with 95% confidence intervals (Bland & Altman, 1986) and by computing the Intraclass Correlation Coefficient (ICC) (Bartko, 1966). In addition, absolute and relative volumetric errors between manual and automatic segmentations were computed for each scan. Absolute error was defined as the voxel-wise difference in corpus callosum volume between methods, and relative error as the percentage difference with respect to the manual reference. These metrics provide an intuitive measure of how closely automatic segmentations reproduce manual delineations.

Because volumetric agreement alone does not ensure spatial fidelity of the segmentation, spatial similarity metrics were also computed for each pair of masks. The Dice Similarity Coefficient (DSC) (Zheng et al., 2022) quantified spatial overlap, while the Hausdorff Distance (HD) (Battalapalli et al., 2022) measured the maximum boundary discrepancy between segmentations in millimeters, providing complementary information regarding contour accuracy.

To stratify animals according to phenotype, a qualitative assessment of the corpus callosum volume in the T1 scans was performed. This validation was carried out with 7 scans classified as controls and 7 scans classified as hypoplastic.

Data were treated as normally distributed based on visual inspection of distributions and sample size considerations. All statistical analyses and plotting were conducted using R (version 4.4.2). Spatial similarity metrics (DSC and HD) were computed in Python using a Dockerized local instance of Google Colab.

## Results

Thirty original marmoset scans were used to validate the automatically generated segmentation masks. The mean volumes obtained from manual delineations (201 voxels) and automatic segmentations (205 voxels) did not differ significantly (paired t-test, p = 0.216). The mean difference corresponded to approximately 2% of the average corpus callosum volume, measured in number of voxels (Fig. 3A).

**Figure 3.**
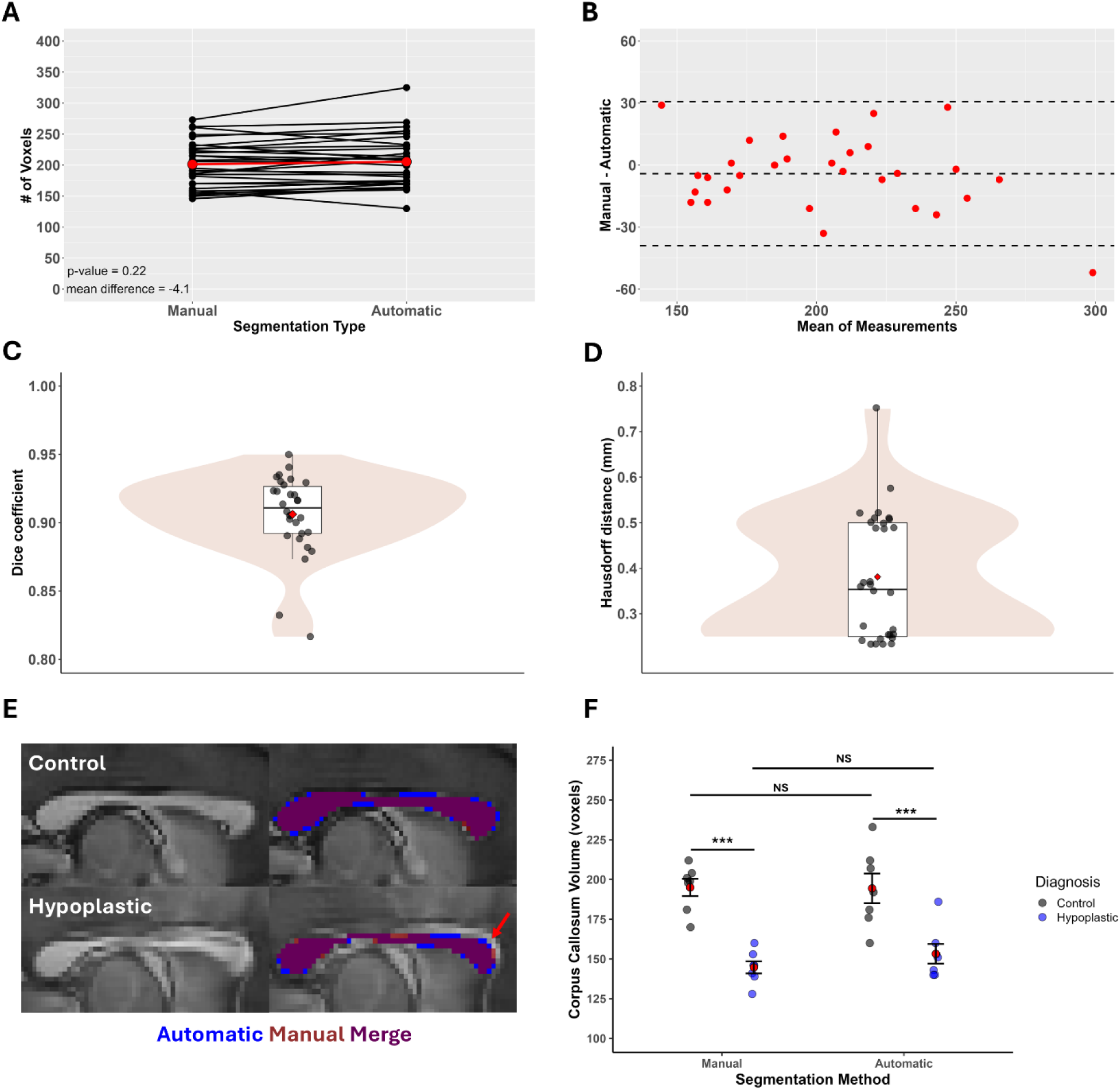
Validation of the automatically segmented masks in marmoset scans. (**A**) Scatterplot showing the distribution of volumes from 30 manually delineated and automatically generated masks. No statistically significant differences were detected. (**B**) Bland-Altman plot displaying the difference between measurements on the y-axis and the mean of measurements on the x-axis. No systematic bias was observed. (**C**) Dice Similarity Coefficient demonstrating strong volumetric agreement between manual and automatic segmentations. (**D**) Hausdorff Distance indicating submillimetric contour differences between segmentations. (**E**) Left: representative examples of corpus callosum morphology in control and hypoplastic animals. Right: color overlays showing manual and automatic masks, and their overlap. (**F**) Differences in corpus callosum volumes between control and hypoplastic animals detected using both segmentation methods (*** p < 0.001).

Bland-Altman analysis revealed no systematic bias between the two methods. The mean difference line was close to zero (showing that our method was not biased), with all data points lying within the 95% confidence interval except one, and no trend was observed as the magnitude of the measurements increased (Fig. 3B). Agreement analysis demonstrated excellent reliability between methods, with an intraclass correlation coefficient (ICC, two-way agreement model) of 0.896 (95% CI: 0.795–0.949).

Spatial similarity metrics further supported the robustness of the model. The average DSC exceeded 0.90, with only two cases presenting values between 0.80 and 0.85 (Fig. 3C). HD was submillimetric in all cases, with a mean value below 0.4 mm / 1.75 voxels (Fig. 3D).

In addition, absolute and relative error analyses demonstrated that the automatic method deviated on average by only 13.7 ± 11.8 voxels from the manual reference, corresponding to a mean percentage error of 6.82 ± 5.46%. These values further indicate that the automatic segmentation closely reproduces manual delineations with minimal volumetric deviation.

Figure 3E illustrates corpus callosum morphology in two different phenotypes: control and hypoplastic. In the right panel, we show a high degree of overlap between manually delineated and automatically generated masks. In the hypoplastic example, the model accurately segmented the corpus callosum excluding the potential contamination of the adjacent pericallosal artery (red arrow). As observed with the manually drawn masks (p = 0.0001), volumetric differences between control and hypoplastic animals remained detectable when using the model-generated masks (p = 0.0009).

## Discussion

Brain malformation comprises a highly heterogeneous group of conditions with varied anatomical presentations (Thalhammer et al., 2025), making the accurate delineation of specific brain structures a challenging task both in neuroimaging studies and clinical settings. In this context, segmentation methods must be robust to anatomical variably while remaining reliable in scenarios where large training datasets are not available. To our knowledge, this is the first study to implement an imaging augmentation pipeline capable of introducing robust, yet anatomically plausible variations specifically designed for segmentation tasks in the context of structural brain malformations. Because deep learning segmentation models rely on learning features across varying presentations, this pipeline substantially reduces the dependance on large datasets from real subjects – a common limitation when studying rare conditions with low clinical prevalence. Rather than relying on dataset size, our approach leverages geometrically induced variability to enrich the feature space available to the model during training.

Our model was validated in the marmoset, as animal models are widely used in neuroscience research, particularly for investigating brain structural development (e.g., of the corpus callosum) (Edwards et al., 2020; Szczupak et al., 2020a; Szczupak et al., 2020b), as well as in preclinical studies of neuropsychiatric disorders (Khanbabaei et al., 2019; Magalhães et al., 2024; Oikonomidis et al., 2017). The common marmoset (Callithrix jacchus) is an ideal model to study neuroscience as it is a New World primate that can bridge the gap between rodents and humans due to its similarities in brain size to the rodent yet maintaining primate-specific evolutionary features, such as their expanded frontal cortex (Schaeffer et al., 2020). Furthermore, the common marmoset reaches sexual maturation at around 12-18 months with a gestation period of about 145 days, allowing for a litter of up to 2-3 pups per year (Sasaki, 2015). This yield is higher compared to an Old-World primate such as the Macaca mulatta that can give birth to one monkey within a gestation period of 146-180 days. This reproductive feature of the marmoset is especially relevant for generating more transgenic primate models within a shorter time frame. Similarly, basic research in transcriptomics has shown that gene expression in neuronal systems differs in primates relative to rodents (Sasaki, 2015).

A central methodological contribution of this work lies in the combination of two complementary augmentation strategies. First, rigid transformations were applied at the volumetric level prior to slice extraction. Although rigid transformations alone do not alter anatomy, when combined with fixed-plane slicing, they generate apparent anatomical variability at the slice level due to changes in the intersection angle between the slicing grid and the rotated volume. Second, additional in-plane slice augmentations were applied during training. These two steps operate in different geometric domains and together create a training set that exposes the model to a wide range of plausible anatomical presentations without requiring additional subjects.

We designed a multi-layered validation framework to ensure that the model’s performance was not evaluated solely on volumetric agreement. Paired comparisons between manually and automatically generated masks revealed no statistical differences, with a very small absolute mean difference. Bland-Altman and ICC analyses demonstrated strong agreement and absence of systematic bias between methods. Importantly, spatial similarity metrics further confirmed segmentation fidelity: the average Dice Similarity Coefficient exceeded 0.9, while the Hausdorff Distance remained submillimetric. These results indicate not only a high voxel overlap but also high contour accuracy between manual and automatic segmentations.

To further test the practical utility of the model, we stratified animals according to corpus callosum phenotype and assessed whether known volumetric differences between groups could still be detected using the automatically generated masks. The model preserved group differences with comparable effect sizes, demonstrating that it does not merely reproduce volumes but maintains the biological signal necessary for downstream analyses. This is particularly relevant in studies of hypoplasia, where differences are subtle and may not be readily identifiable through qualitative inspection. Corpus callosum dysgenesis encompasses, in general, three different clinical presentations, and although, two of these are easily detectable via qualitative assessment of MRI scans - agenesis (the complete absence of the structure) and partial dysgenesis (when the corpus callosum is partially formed) - the model is able to detect hypoplasia, when the corpus callosum is fully formed and the difference is at the volume level, which might not be easily perceived in a qualitative assessment.

Beyond accuracy, the speed of segmentation represents a significant practical advantage. While manual delineation is feasible for small datasets, it becomes impractical as sample sizes grow. The model’s ability to generate masks in seconds (1 mask per second) enables large-scale quantitative studies that would otherwise be prohibitively time-consuming.

## Conclusion

The present work introduces a methodological framework that enables the training of robust segmentation models for anatomically variable brain structures using a limited number of original scans. By combining whole-volume rigid transformations with slice-level augmentation, we demonstrate that it is possible to generate sufficient anatomical diversity to train a highly accurate U-Net model for corpus callosum segmentation in the marmoset. Importantly, this framework is not restricted to the corpus callosum nor to this species. Rather, it establishes a translational strategy for studying rare brain malformations, where manually delineating large datasets is often impractical. The marmoset serves here as an ideal experimental model in which controlled anatomical variability can be leveraged to develop and validate segmentation tools that are conceptually transferable to human neuroimaging contexts.

From a clinical perspective, the corpus callosum is among the structures most frequently affected in developmental and neurological disorders. A tool capable of rapidly and reliably estimating its volume has direct implications for both research and diagnostic workflows. Such automated segmentation can support the extraction of normative values, assist radiologists in identifying subtle hypoplastic presentations that might escape qualitative inspection, and enable large-scale morphometric studies that would otherwise be unfeasible due to the burden of manual annotation.

Therefore, beyond providing an accurate segmentation model, this study proposes a generalizable augmentation-driven training strategy that bridges experimental neuroscience models and clinically relevant neuroimaging applications.

## Limitations and perspectives

Despite the robustness of the proposed augmentation pipeline, it primarily addresses anatomical variability and does not yet incorporate variability related to acquisition protocols or scanner-dependent artifacts. The inclusion of texture-based augmentation methods, such as Perlin noise (Bae et al., 2018) and Gaussian noise injection (Khalifa et al., 2022) could substantially enhance the generalizability of the trained networks (Akbiyik, 2023; Goceri, 2023). Although the framework is modality-agnostic in principle, it has so far been validated exclusively on T1-weighted images. Future work should investigate its applicability across other MRI contrasts and acquisition settings.

Another important perspective is the extension of this approach to other types of central nervous system malformations. Conditions such as congenital Zika virus syndrome, which present far more severe and complex anatomical disruptions (de Castro et al., 2017; Vhp et al., 2020) than hypoplasia, constitute an ideal scenario to test the limits of augmentation-driven training strategies. In such contexts, the ability to synthetically expand anatomically plausible training datasets may be decisive for enabling reliable automated segmentation.

An additional direction for future work is to systematically evaluate hybrid training strategies that combine original scans with augmented data. Training exclusively on synthetic or augmented images can increase sample diversity and reduce dependence on large real-world datasets, but it may also introduce a domain gap if the transformed images do not fully capture the variability present in real acquisitions. Incorporating real scans into the training set may improve biological and acquisition-level fidelity, whereas augmented samples may continue to provide the diversity needed to prevent overfitting in small cohorts. A useful next step would therefore be to train and compare models under different real-to-augmented ratios, in order to identify an optimal balance between realism and variability for robust performance in real-world cases.

Finally, while the current model already operates at high speed during inference, the training process remains computationally demanding. Ongoing efforts aim to reduce network complexity and optimize training efficiency without compromising segmentation accuracy. Together, these perspectives reinforce that the present tool should be seen not as a finalized solution, but as a foundational step toward a broader translational framework for studying brain malformations through automated image analysis.

## Acknowledgements

The Article Processing Charge (APC) for the publication of this research was funded by the Coordenação de Aperfeiçoamento de Pessoa de Nível Superior (CAPES) (ROR identifier: 00x0ma614). For the purpose of Open Access, the authors have applied a Creative Commons Attribution (CC BY) license to any Author Accepted Manuscript (AAM) version arising from this submission.

The authors acknowledge the use of ChatGPT (OpenAI) to improve the readability and language style of this manuscript. All AI-generated text was carefully reviewed and edited by the authors, who take full responsibility for the final content of the manuscript.

The authors declare that they have no competing interests.

